# Prediction of biological age by morphological staging of sarcopenia in *Caenorhabditis elegans*

**DOI:** 10.1101/2021.06.16.448702

**Authors:** Ineke Dhondt, Clara Verschuuren, Aleksandra Zečić, Tim Loier, Bart P. Braeckman, Winnok H. De Vos

## Abstract

Sarcopenia encompasses a progressive decline in allover muscle quantity and quality. Given its close association with aging, it may represent a valuable healthspan marker. Given the strong commonalities with human muscle structure and the facile visualization possibilities, *C. elegans* represents an attractive model for studying the relationship between sarcopenia and healthspan. However, classical assessment relies on visual scoring of muscle architecture, which is subjective and inaccurate. To resolve this, we have developed an automated image analysis pipeline for the detailed quantification and classification of muscle integrity in confocal microscopy images from a cohort of aging myosin::GFP reporter strains. We then extracted a variety of morphological descriptors and found a subset to scale linearly with age. This allowed us to establish a general linear model that predicts biological age from a morphological muscle signature. To validate the model, we evaluated muscle architecture in long-lived worms that are known to experience delayed sarcopenia by targeted RNAi-mediated knockdown of the *daf-2* gene. We conclude that quantitative microscopy allows for staging sarcopenia in *C. elegans* and will be of use for systematic screening for pharmacological or genetic modulators that mitigate age-related muscle frailty and thus improve healthspan in *C. elegans*.

## Introduction

Aging is a complex phenomenon, which can be defined as a progressive deterioration of cell and tissue functions in living organisms with age (Gems, 2014). For long, biogerontologists exclusively focused on the determinants of lifespan with an eye on its extension in diverse organisms. However, rather than just extending lifespan *per se*, it is becoming increasingly clear that it is more desirable to improve the quality of life by limiting the risk for frailty and disease at advanced age (Olshansky, 2018). Hence, extending healthspan, the life period in which one is functionally independent and free from serious disease, has now become a central theme of modern biogerontology (Luyten et al., 2016; Rattan, 2018). A key component of human late-life frailty is sarcopenia, a progressive decline in muscle quantity and quality (Evans et al., 2010; Fulop et al., 2010). A delayed onset of sarcopenia may thus represent a valuable biomarker of extended healthspan. Due to its genetic amenability and short lifespan, *C. elegans* has become an invaluable model organism to the study of aging. As its soma is post-mitotic and transparent, the progression of several age-related pathologies can easily be followed *in vivo*. Aging *Caenorhabditis elegans* experience gradual, progressive muscle deterioration, resembling human sarcopenia. In young nematodes, body wall muscle myofilaments are well-organized in a tight, parallel manner, while in older animals, they show progressive disorganization, irregular orientation, breaks, and abnormal bends (Herndon et al., 2002). The ease with which muscle structure can be visualized *in vivo* and the similarity of its muscle microanatomy to that of man make *C. elegans* a strong model for studying sarcopenia (Christian and Benian, 2020; Herndon et al., 2002; Meissner et al., 2009). Additionally, the lack of muscle stem cells in this nematode model allows focusing on muscle deterioration during aging without the confounding influence of muscle regeneration (Christian and Benian, 2020). Worm myofilament organization is often studied by visual assessment of a reporter strain expressing fluorescent markers such as GFP-tagged myosin heavy chain (MHC) A in the body wall muscle cells (Campagnola et al., 2002; Glenn et al., 2004). Former studies relied almost exclusively on manual scoring of myofilament organization and the appearance of muscle defects (Ben-Zvi et al., 2009; Brehme et al., 2014; Glenn et al., 2004; Herndon et al., 2002; Karady et al., 2013; Lamarche et al., 2018; Meissner et al., 2009; Meissner et al., 2011). Such quantifications are prone to observer bias and are not very sensitive. Hence, we developed an objective analysis tool for the detailed quantification of muscle integrity in myosin::GFP reporter strains. In order to stage the severity of sarcopenia in aging muscle cells, we extracted a variety of morphological descriptors (features) of individual myofilaments within the muscle cell using an automated image-processing pipeline. We found that a subset of features scaled linearly with age, allowing us to establish a linear model to predict biological age from a morphological muscle signature. To prove its potential for screening, we validated our model on a biologically relevant mutant displaying delayed sarcopenia (Depuydt et al., 2013; Herndon et al., 2002; Wang et al., 2019).

## Materials and Methods

### C. elegans maintenance

In this study, we used the transgenic strain RW1596 *stEx30* [*myo-3p::gfp::myo-3 + rol-6(su1006)*]. Animals were maintained on nutrient agar plates with *E. coli* K12 prior to experiments. NGM plates containing Agar NO.1 (2.5 % w/v, Oxoid), Pepton N-Z-Soy(R) BL4 (0.25% w/v, Sigma Aldrich), NaCl (0.3% w/v), cholesterol (0.0005% v/v), CaCl_2_ (1 mM), MgSO_4_ (1 mM), K_2_HPO_4_/KH_2_PO_4_ (25 mM, pH 6.0), carbenicillin disodium (0.5 mg/ml, Fisher BioReagents™) and Isopropyl-β-D-thiogalactopyranoside (IPTG Dioxane-free, 1 mM, Fisher BioReagents™) were used for experiments. As future muscle integrity studies will rely mostly on the use of RNAi, we used the bacterial control strain *E. coli* HT115 containing the (empty) L4440 vector as food source. Bacteria were grown overnight on a shaker (120 rpm) at 37°C in LB Broth Lennox (2% w/v, Fisher Bioreagents) containing carbenicillin disodium (2 mg/ml, Fisher BioReagents™). IPTG (1 mM, Fisher BioReagents™) was added the next day and the culture was placed on a shaker (120 rpm) for an additional 2 hours at 37°C. Bacteria were washed with salt water (0.3 % NaCl w/v) and concentrated five times. Seeded NGM plates were kept at 20°C to allow bacterial growth overnight. Synchronized L1 nematodes were spotted onto these plates and FUDR (100 μM, Acros Organics) was added at L4 stage to prevent offspring. Animals were transferred regularly to fresh plates during the experiment.

### Image acquisition

A Nikon TiE-C2 confocal laser scanning microscope, equipped with a 40xWI Plan Apo objective (NA 1.20, water immersion) was used for imaging. Images (1024 × 512 pixels, 6.67 pixels per μM, scanner zoom 2x) were collected by exciting with a 488-nm solid state argon laser and detecting through a 525/50 nm bandpass filter. Laser power (1.25%), gain (82) and offset (5) were maintained constant throughout all experiments. Images were collected for 4 independent aging cohorts at specific time-points during aging (at day 1, 4, 6, 8, 12, 14 and 18 of adulthood). Standardized image nomenclature was used to facilitate downstream analysis. We recommend using the following format: C01W01M01.nd2, in which C, W and M refers to the Condition, Worm and Myofilament, respectively. The total number of worms and muscle cells per condition are summarized in Table 1.

**Table 1.** Number of observed animals and recorded muscle cell spindles (between brackets) are indicated for each time point per studied cohort.

|  | Aging cohort |  |  |  |  |  |  | Validation (day 14) |  |
| --- | --- | --- | --- | --- | --- | --- | --- | --- | --- |
|  | day 1 | day 4 | day 6 | day 8 | day 12 | day 14 | day 18 | EV | *daf-2* |
| **Cohort** | 10 | 10 | 10 | 10 | 10 | 5 (12) | NA | 11 (55) | 12 (59) |
| **Cohort** | 9 (16) | 5 (11) | 10 | 7 (19) | 7 (17) | 9 (24) | 10 | 10 (45) | 12 (53) |
| **Cohort** | 6 (14) | 9 (19) | 9 (22) | 11 | NA | 7 (17) | 12 | 10 (48) | 15 (50) |
| **Cohort** | NA | 8 (14) | 10 | NA | NA | 9 (23) | 9 (22) | NA | NA |
| **Total** | 25 | 32 | 39 | 28 | 17 | 30 | 31 | 31 (148) | 39 (162) |

### Manual scoring of muscle deterioration

Images of body wall muscle cells were evaluated in a randomized and blinded manner using Blinder freeware (Cothren et al., 2018). Each cell was assigned one of three qualitative classes of muscle deterioration: (1) normal, healthy cell with well-organized myofilaments, (2) moderately disorganized cell and (3) severely distorted cell.

### Image analysis

All image analysis was performed in FIJI open-source image processing software (Schindelin et al., 2012). A macro script (musclemetrics.ijm) was written for the assessment of muscle organization in *myo-3*::GFP worms, which is available upon reasonable request. The script consists of a set of interactive tools for cropping and aligning individual muscle cells as well as an automated routine for segmentation and morphological feature extraction of individual myofilaments (Fig. 1). In brief, the algorithm consists of local contrast enhancement, background subtraction and a directional second order edge enhancement (using FeatureJ plugin; https://imagescience.org/meijering/software/featurej/), followed by binarization according to Yen’s autothresholding algorithm. Subsequent particle analysis is done with exclusion of single pixel objects so as to remove noise contributions. Apart from the morphological and intensity metrics that are extracted by default in ImageJ, the analysis also includes a measurement of local thickness and curvature variations. Local thickness descriptors (average, range, variation) are directly based on the output of the Local Thickness plugin (https://imagej.net/Local_Thickness). Curvature is expressed as a deviation of the object’s perimeter with respect to that of a reconstructed version based on a subset of (first 5) elliptic Fourier descriptors using the Fourier Shape Analysis plugin (https://imagejdocu.tudor.lu/plugin/analysis/fourier_shape_analysis). The inverse ratio of these will yield lower values for more irregular objects. Skeletonization of the binary objects provides additional info on the number of branch points and straightness (Feret/Area of the skeleton), whereas the smallest eigenvalues of the structure tensor (as obtained from FeatureJ) are used for calculating the bending energy and determining the number of bend points (local maxima). Finally, a global analysis of myofilament orientation is performed using the Directionality plugin (https://imagej.net/Directionality.html). Per muscle cell, images are stored along with the corresponding regions of interest (ROI) sets and result files.

**Figure 1.**
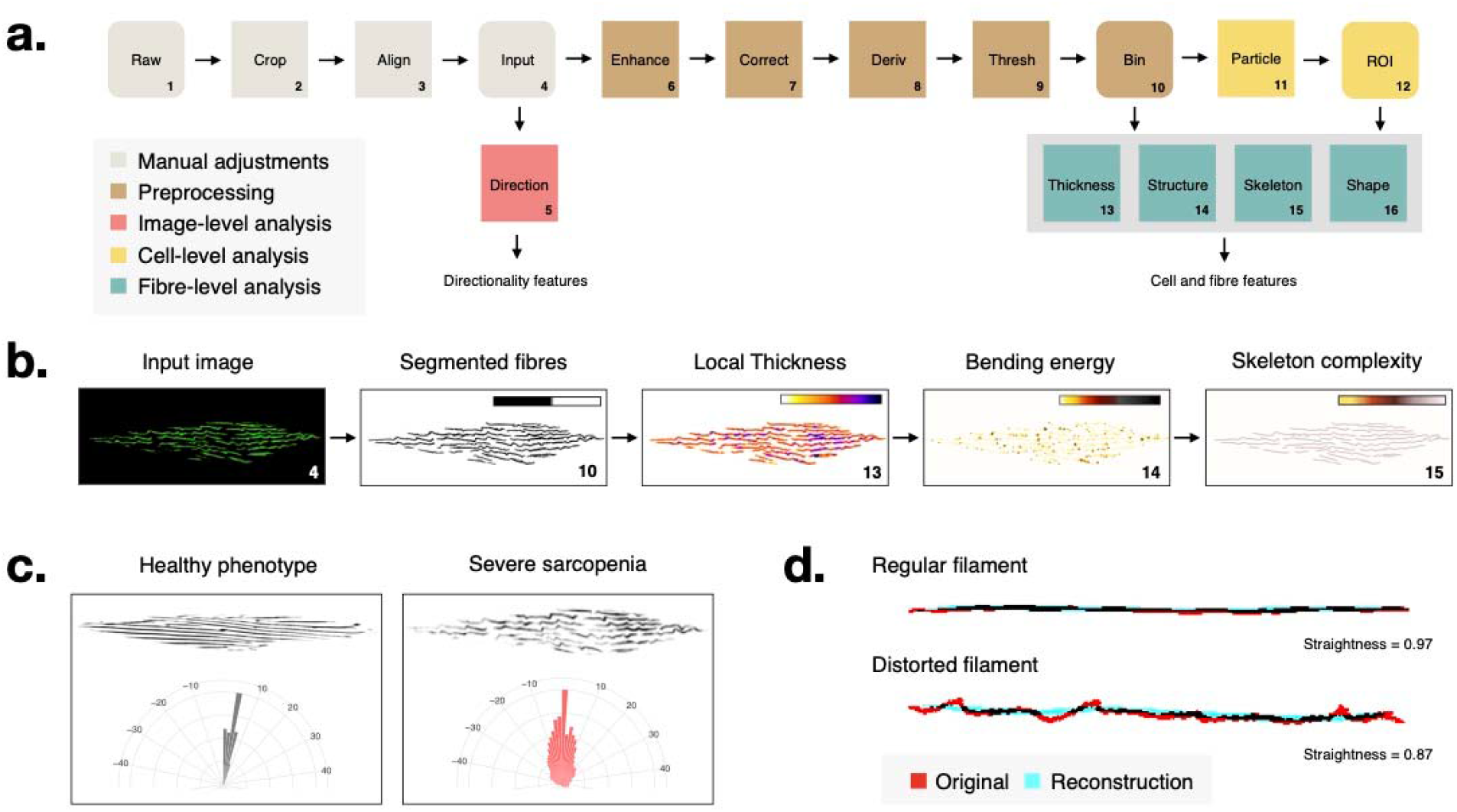
(a) Automated extraction of myofilament (organisation) features using a custom-designed image-processing pipeline, consisting of manual pre-adjustment of the image (cropping and aligning individual cells from raw images resulting in the input image), pre-processing (local contrast enhancement, background correction, directional derivative and thresholding resulting in a binary image) and subsequent image analysis at different levels. The unprocessed input image is used for directionality analysis at the image level, the binary (segmented) image is used for detection of local thickness and structural variations as well as for skeletonization (complexity); and after individual particle analysis, resulting regions of interest (ROI) are used for individual fibre measurements; (b) Illustration of interim results for an image of a severe sarcopenia phenotype (numbers correspond with steps in a); (c) Directional analysis in a healthy vs. severely affected muscle cell. The orientation of myofilaments in the latter deviates significantly from the expected tight parallel myofilament organization observed in young animals, which is reflected in a broader distribution of orientation angles on a radar plot; (d) Fibre straightness is measured as the ratio of the perimeter of the original object (fibre) versus a reconstruction from a limited number of Fourier shape descriptors.

### Statistics

All data analysis was performed in R Studio, using R version 4.0.2. Plotting was done using the ggplot2 package and summary statistics were calculated using the ddply package. All individual result files were loaded into a single data frame, collated with the appropriate metadata regarding experimental conditions (including factors such as age, worm, muscle, and replicate). During image analysis, features were extracted at the individual myofilament level. To evaluate changes at the cellular level, the average as well as the covariance of these values were calculated per muscle cell. These secondary data were combined with features that are inherently derived on a per cell basis such as directionality, global cell size and the total number of myofilaments. Linear models were established using the native lm functions and collinearity checks were performed using the variance inflation (VIF) inspection tool of the car package as well as the correlation detection of the caret package. General linear models with smoothing terms were established using the mgcv package. Statistical comparisons were done using Welch T-test after having verified for normality (Shapiro test and qq-plot) and homoscedasticity (Levene’s test).

## Results

### Sarcopenia occurs progressively with age in C. elegans

Muscle deterioration is a common pathology observed in old animals (Christian and Benian, 2020). To document its penetrance and evolution in *C. elegans*, we set out to score the muscle phenotype with increasing age. To evaluate muscle architecture *in vivo*, we used a transgenic *myo-3*::GFP reporter strain, which expresses GFP-tagged myosin heavy chain (MHC) A in the body wall muscle cells. Confocal images were collected throughout the adult lifespan (day 1 till day 18) of four independent worm populations. At set time points, several muscle cells (n_muscle_ = 2-4) of multiple worms (n_worm_ = 5-12) were visualized using consistent acquisition settings (Table 1). As expected, and in line with earlier findings (Herndon et al., 2002), we observed a marked change in the organization of muscle cells with advancing age, indicative of ensuing sarcopenia. Young muscle cells typically displayed a tightly organized architecture of parallel oriented, thick myofilaments, whereas muscle cells of later life stages (day 8 till day 18), more frequently displayed bends or breaks in individual myofibers leading to a less aligned pattern (Fig. 2a). To gain a first qualitative impression of the age-dependence, we manually attributed a score to the muscle phenotype based on a visual assessment. We thereby discriminated three phenotypes, representing either a healthy well-organized cell, a moderately disorganized cell, or severely distorted cell. To avoid bias, scores were assigned in a blinded manner. While the majority of muscle cells indeed showed a normal, healthy phenotype (75%) at day 1, the fraction of moderately or severely disordered cells progressively increased with age (Fig. 2b). At day 14 and 18, approximately 75% of the muscle cells was found at least moderately distorted. This shows that the degree of sarcopenia increases gradually with age and suggests that its severity may serve as a biomarker for biological age.

**Figure 2.**
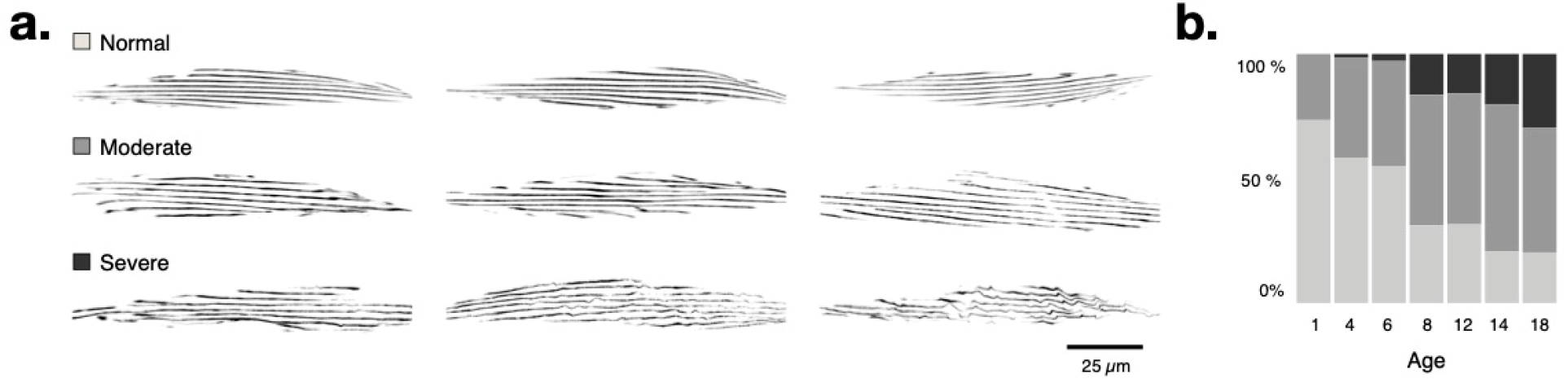
Age-dependent defects in structural organization of the C. elegans body wall muscle. Representative confocal images of individual *C. elegans* muscle cells according to three qualitative classes of muscle deterioration; (b) Blinded, manual scoring of randomized images shows the progressive loss of myofilament organization with age as evidenced by the decrease in normal and increase in moderate to severe phenotypes.

### Automated analysis reveals changing myofilament structure and heterogeneity with age

Having established a strong correlation between age and the number of cells with distorted filaments, we next asked whether we could describe muscle organization more quantitatively so as to enable an objective, automated assessment. To this end, we developed an image analysis routine that extracts a variety of morphological descriptors (features) of individual muscle cells and their constituent myofilaments (Fig. 1, see Materials and Methods section). When inspecting the extracted feature space as a function of the manually attributed score, a number of clear trends could be observed (Fig. 3a, b; Suppl. Fig. S1). Specifically, the fraction of smaller, more compact myofibers increased with severity score as indicated by the lower values for average area and perimeter and larger values for average circularity and roundness. This may reflect the progressive myofiber fragmentation that accompanies sarcopenia. At the same time, the number of irregularly shaped, non-linear filaments grew with severity as indicated by the change in average curvature, straightness and bending energy. Not only the average value but also the variability of certain features varied with the attributed score. For example, the covariance of the filament thickness, area, bending energy and solidity increased with severity score, suggesting a larger fiber heterogeneity in more distorted cells. Finally, the orientation of individual fibers became more heterogeneous with sarcopenia score as evidenced by the larger dispersion and lower amount of directionality information that could be explained by Gaussian fit on the dominant orientation. When plotting the exact same features as a function of age instead of score (Fig. 3c; Suppl. Fig. S2a), very similar trends were observed suggesting that these metrics also portray the penetrance of sarcopenia in a population of aging worms.

**Figure 3.**
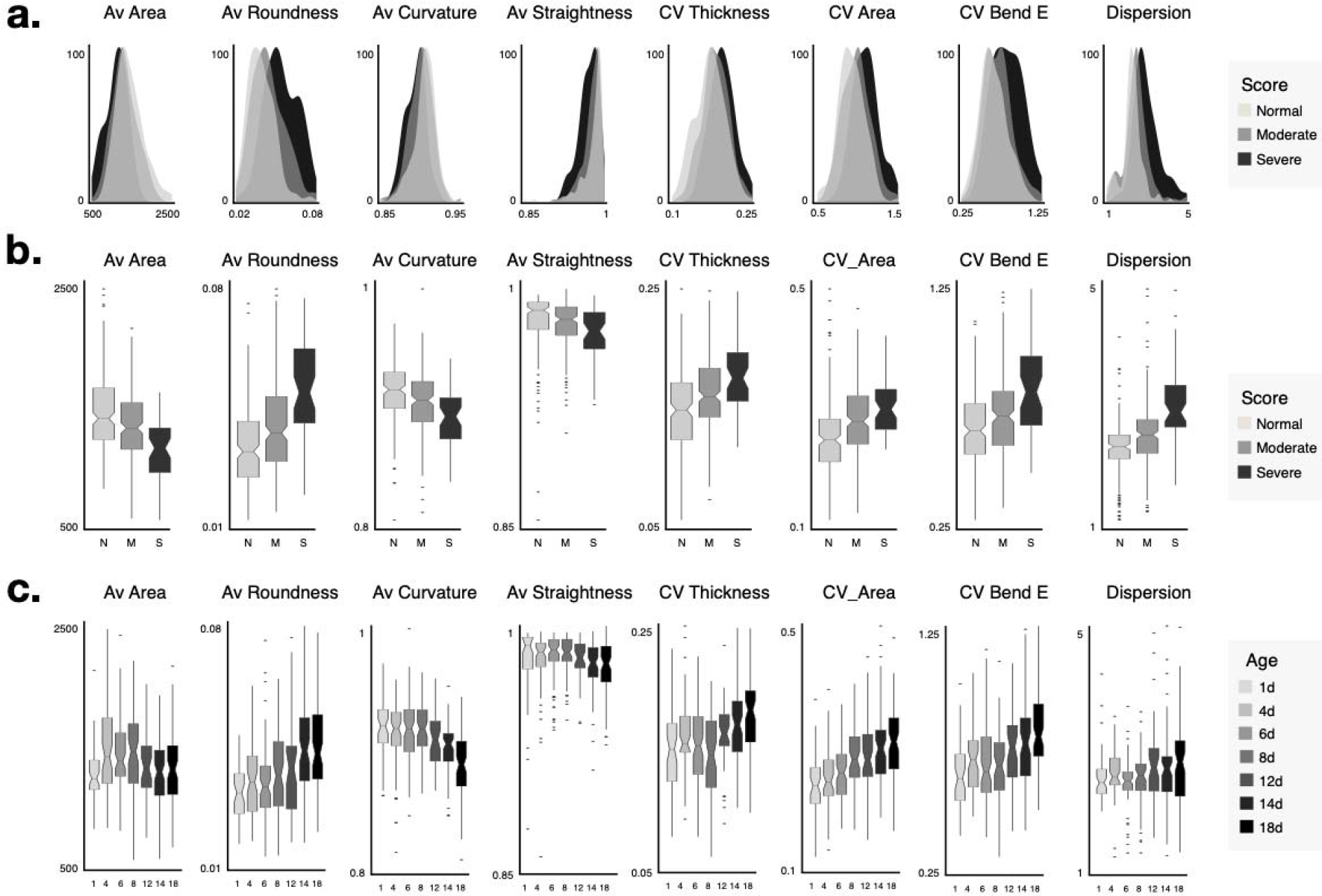
Selected features describing morphological changes as a function of score and age. (a) histograms illustrating feature shifts and variability as a function of score per muscle cell; (b) boxplots of the same parameters per score; (c) boxplots per age show similar trends. (Av = average; CV = coefficient of variation).

### A morphological signature can be used to predict biological age

Given the close resemblance between the feature distributions of severity score and organismal age, we hypothesized that the morphological features could inform on the biological age by providing an estimate of the degree of sarcopenia. In order to confirm this hypothesis, we first statistically queried the age dependence of the individual metrics. Univariate linear regression revealed that the magnitude of many features significantly scaled with age (46/60 features having a p-value < 0.05, Suppl. Fig. S2b), thus confirming the observed trends. However, the Pearson correlation coefficient (R) of individual comparisons was rather low, with a maximum value of R = 0.41 for the average fiber number. We reasoned that a more inclusive model would perform better and therefore, we performed linear regression using all standardized features. The resulting linear model was able to account for a large proportion of the variability as evidenced by an R of 0.72 (adj. R^2^ = 0.45), despite the persistence of normal muscle fibers in older organisms (Fig. 4a).

**Figure 4.**
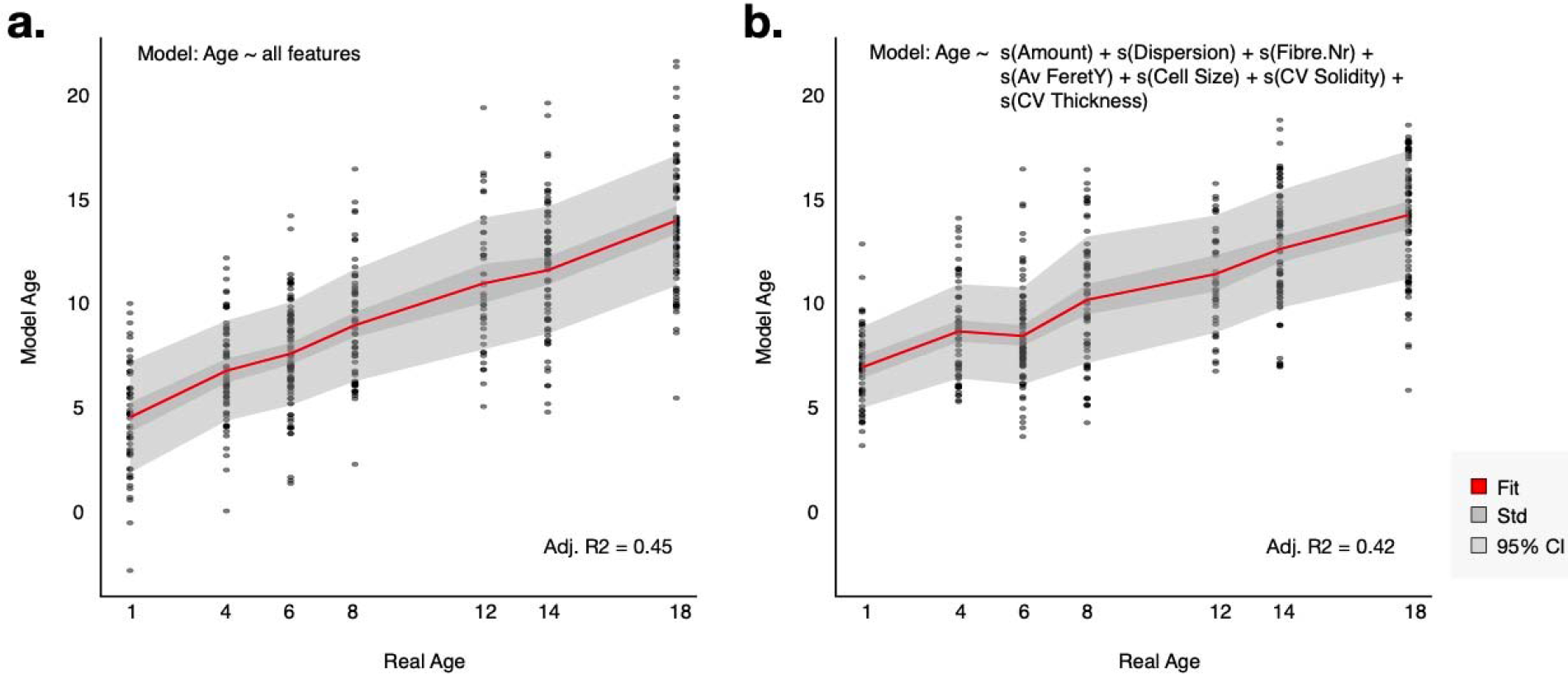
(a) Linear regression on the complete dataset using all features; (b) General additive model with smoothing terms shows comparable performance on the complete dataset.

**Figure 5.**
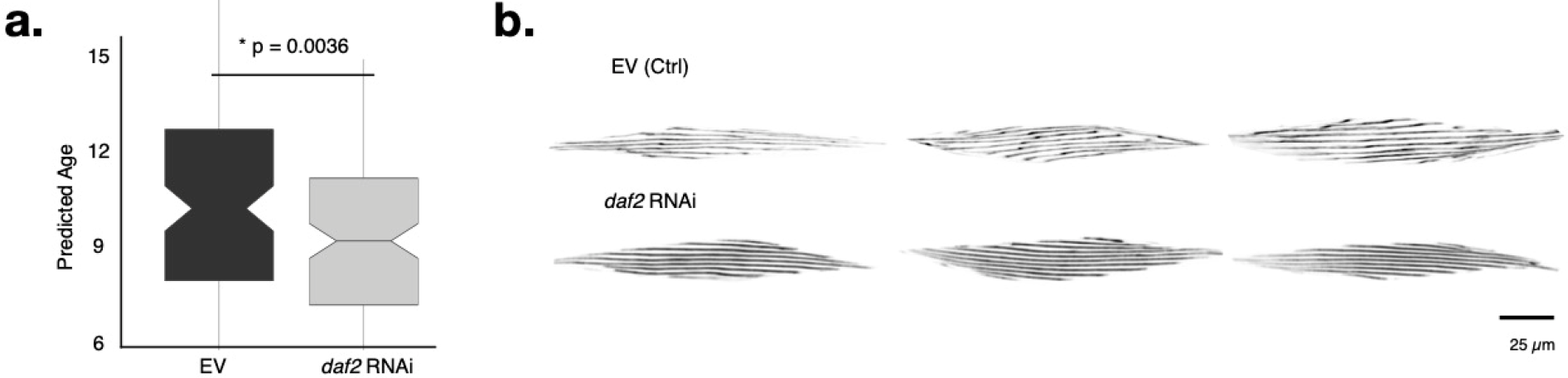
Downregulation of *daf-2* using RNAi leads to a younger biological age at a chronological age of day 14 (a) as predicted by the established GAM; (b) This is also apparent as a more parallel and regular myofilament organisation in muscles of *daf-2* RNAi-treated vs control worms.

Supported by these results, we wondered whether such a model could be used to predict age. To this end, we split our dataset into a training (2/3) and validation (1/3) set and established a linear model on the first to evaluate its performance on the latter. This revealed a comparable performance for the full model (adj. R^2^ = 0.43; RSE = 4.4). Although the most complete model (using all features) will explain the largest proportion of the variability in the training set, it might not necessarily perform best on a separate validation set as it will tend to be biased (overfitted) towards the former. This goes at the expense of its robustness. To establish the most parsimonious model that performs well on both datasets, we narrowed its feature space by setting cut-offs on multi-collinearity, either by removing variables with more than 70% correlation or variance inflation factors above 5 (Suppl. Fig. S3). In doing so, we obtained a model that only included 7 parameters (Average Amount, Average Dispersion, Average Fibre Number, Average Feret Y, CoV Solidity, Average Cell.Size, CoV_Thickness) and performed equally well on the validation set as the full model (adj. R^2^ = 0.46; RSE = 4.2). Introduction of smoothing terms (in a general additive model) slightly improved this further (adj. R^2^ = 0.47; RSE = 4.2). Thus, we conclude that a minimal signature of 7 morphological descriptors, covering cell size, fibre directionality and fibre shape heterogeneity, can be used for estimating biological age in *C. elegans*.

### Morphological staging confirms sarcopenia delay in worms with reduced insulin/IGF1-like signaling

In a final step, we sought to benchmark the approach for predicting biological age in a physiologically validated model. Since reduced insulin/IGF-1 signaling leads to lifespan extension in worms (Kenyon et al., 1993) and hypomorphic mutants of the insulin/IGF-1-like receptor *daf-2* show increased healthspan (Podshivalova et al., 2017) including a delay in the onset of sarcopenia (Kashyap et al., 2012), we decided to use this model. We compared *daf-2* RNAi-treated *myo-3*::GFP worms with empty vector (EV)-treated controls at advanced age (14 day) in three independent biological replicates. The animals were subjected to an identical image acquisition and analysis procedure as before and the resulting standardized feature set was used as input for the established general linear model. Using the model, we observed a significant reduction of the predicted biological age for *daf-2* RNAi worms compared to their control counterparts (P-value = 0.0036, Welch two-sample T-test). Hence, using a biological control, we provided an independent validation of the predictive value of the linear model for sarcopenia staging.

## Discussion

In this work, we focused on the gradual microfilament deterioration in the *C. elegans* body wall muscle, an age-associated pathology that resembles human sarcopenia (Ben-Zvi et al., 2009; Christian and Benian, 2020; Herndon et al., 2002). Although worms only live two to three weeks, their body wall muscles showed clear structural disorder with increasing age. This vulnerability likely originates from the ‘built-for-life’ property of *C. elegans* muscle cells: they are non-replaceable and post-mitotic, while their myofilament proteins turn over extremely slow (Dhondt et al., 2016; Dhondt et al., 2017). While several studies have scored muscle defects in *C. elegans*, the large majority relied on manual scoring (Ben-Zvi et al., 2009; Brehme et al., 2014; Glenn et al., 2004; Herndon et al., 2002; Karady et al., 2013; Lamarche et al., 2018; Meissner et al., 2009; Meissner et al., 2011). Only in one recent study, muscle cell area and myofilament length were measured by image analysis to more objectively evaluate *C. elegans* body wall muscle integrity (Soh et al., 2020). However, this study focused on mutants with severe muscle defects and while these two parameters may have had sufficient power to discriminate overt phenotypes, they do not have the resolution or sensitivity to capture the full extent of gradual morphological changes that take place during sarcopenia development. Hence, we introduced an unbiased approach to stage sarcopenia.

Having confirmed that the phenomenon is highly age-dependent, we reasoned that its quantitative description could serve as a predictor of biological age and *in extensu* of healthspan, comparable to other *C. elegans* healthspan biomarkers such as those related to muscle function and stress resistance (Hahm et al., 2015; Keith et al., 2014). Starting from an exhaustive extraction of morphological features from segmented muscle cells, a non-redundant set was retained that, together, described best the changes that are observed in muscle cells with biological age. Their biological relevance can be explained and, in most cases, confirms earlier microscopic observations. Indeed, Herndon *et al*. (Herndon et al., 2002) found that aged worms have a loss of direction of individual sarcomeres and the stochastic nature of worm sarcopenia described in that same paper is in accordance with the age-related increase in covariance we observed in several features. Of note, increased cellular heterogeneity appears to be a common feature in aging as it was also established in a study of biophysical and molecular properties of human fibroblasts from an aging cohort (Phillip et al., 2017). Myofilament breaks and bends are also often visible in old muscle cells and explain the smaller and more detected fibre entities. The increase in allover cell size is at odds with the notion that sarcopenia is characterized by reduction in muscle mass and concomitant flattening of the body wall muscle (Herndon et al., 2002).

Of note, a significant proportion of the *C. elegans* muscle cells retain a healthy phenotype at very advanced age. Although genotype and environment were held constant (within and between aging cohorts), we observed large cell-to-cell variation in the degree of muscle deterioration within individuals, but also considerable inter-individual variability. This variability was noted earlier as well (Herndon et al., 2002) and contributed to the uncertainty of our model. Notwithstanding the high number of ostensibly normal cells, the model was able to classify and predict biological age. While increasing the number of sampled animals would most likely improve its accuracy, this is not easily manageable with a standard setting using individual microscopy slides. Hence, leveraging the approach to a high-throughput setting would demand a more controlled sampling, *e.g.,* by organism-on-chip microscopy (Cáceres et al., 2018; Kim et al., 2020; San-Miguel and Lu, 2013). Ideally, this could be combined with automated detection and delineation of muscle cells, which would make time-consuming manual segmentation steps unnecessary. However, due to unclear outlines of body wall muscle cells, this would require more advanced detection algorithms, *e.g.,* based on deep learning, than the currently available pipelines. Next to increasing the number of sampled worms to increase prediction accuracy, it may be worth investigating whether the variability in muscle phenotype can be reduced. In old worms, about 25% of the myofilaments did not show obvious deterioration at the light microscopical level, but it is conceivable that sub-threshold, molecular damage may be present that is not readily visible under standard settings. A controlled paradigm to evoke mechanical stress, such as stimulated swimming in a shaking liquid culture over a fixed time period, may elicit the underlying molecular damage and result in a more uniform and outspoken myofilament phenotype in old worms.

In conclusion, we here present a tool that is capable of predicting *C. elegans* healthspan, based on myofilament features, as exemplified by our analysis of an insulin/IGF1-like signaling mutant. In its current form, this tool can be used downstream of the many high-throughput screens for genes, treatments and compounds that modulate healthspan in *C. elegans* (Bulterijs and Braeckman, 2020; Le et al., 2020; Luyten et al., 2016; Sayed et al., 2021). It objectively probes for the effect of such treatments on sarcopenia, an age-related pathology conserved from worms to humans. Further automation and upscaling of the protocol may lead to the first *C. elegans*-based high-throughput sarcopenia screening platform.

## Acknowledgements

We thank the Caenorhabditis Genetic Center (CGC) for providing the RW1596 strain used in this study.

## Ethical guidelines statement

All authors listed on this manuscript have made a substantial contribution to the work and have critically read and approved the final version of the manuscript. The animal work complies with the ethical committee guidelines of the University of Ghent. The authors of this manuscript certify that they comply with the ethical guidelines for authorship and publishing in the Journal of Cachexia, Sarcopenia and Muscle (Haehling et al., 2019).

## Author contributions statement

Data collection: ID, CV, AZ, TM; Software development: WHDV; Data analysis: WHDV, ID; manuscript writing: ID, WHDV, BPB.

